# Developmental divergence in gene regulation among rapidly radiating cichlid species

**DOI:** 10.1101/2024.01.24.577063

**Authors:** Anna Duenser, Pooja Singh, Ehsan Pashay Ahi, Marija Durdevic, Sylvia Schaeffer, Julian Gallaun, Wolfgang Gessl, Ole Seehausen, Christian Sturmbauer

## Abstract

Developmental shifts in gene regulation underlying key innovations remain largely uncharacterized over short evolutionary timescales. Here, we investigate the gene regulatory landscape of trophic innovations in the fastest vertebrate adaptive radiation: cichlid fishes from Lake Victoria. By analyzing the whole-transcriptomes of the oral and pharyngeal jaws from two life stages in five species adapted to divergent trophic niches, we find that both gene and isoform expression are developmentally dynamic. Expression signatures are most similar across jaws at the early stage and then diverge into species-specific developmental programs in adults. However, even at the early stage, expression in the oral jaws of species adapted to herbivory is distinct from those of carnivores, a pattern not observed for the pharyngeal jaws. We further show that differentially expressed and spliced genes between herbivorous and carnivorous species regulate different genes and pathways, particularly in adults. Interestingly, we find that splicing-mediated exonization of a craniofacial development gene, kaznb, may have contributed to the evolution of herbivory in Lake Victoria cichlids. Overall, our results contribute to our understanding of how ontogenetic shifts in gene regulation can facilitate rapid adaptive evolution.

## Introduction

Since King and Wilson (1975) concluded that variation in protein sequence alone cannot explain the phenotypic differences between human and chimpanzees, transcriptional regulation has been shown to play a fundamental role in phenotypic evolution and diversification (Brawand et al. 2011; Barbosa-Morais et al. 2012; Merkin et al. 2012). However, few studies have focused on recently diverged species, where regulatory changes are hypothesised to dominate phenotypic divergence, as they can emerge more rapidly from standing genetic variation and relaxed selection than *de novo* protein-coding variation (Singh et al. 2017; El Taher et al. 2020; Richards et al. 2021; Singh et al 2021; Singh and Ahi 2022; Carruthers et al. 2022). Even fewer studies have looked at regulatory changes during development in recent and rapid bursts of diversification (Cardoso-Moreira et al 2019).

Cases of adaptive radiation, where species diversify rapidly in tight connection with ecology, provide promising model systems to study the role of gene regulation in rapid evolutionary diversification (Abzhanov et al. 2006; Gillespie et al. 2020). The radiations of cichlid fishes in the Great East African lakes are such model systems where rapid diversification proceeded along many ecological and morphological axes (reviewed in Kocher 2004). The Lake Victoria (LV) cichlid radiation comprising ∼500 species is arguably one of the most spectacular of these radiations, having emerged after the complete desiccation of the lake <16,000 years ago (Johnson et al. 2000; Seehausen 2002). The functionally decoupled oral and pharyngeal jaws of cichlids is hypothesised to be a key innovation and basis of their evolutionary success that enabled rapid trophic diversification (Fryer and Iles 1972; Parsons et al. 2011). It was recently found that cichlids dominate the biomass of Lake Victoria not because they arrived first as the lake filled up, but because they had ecologically versatile trophic morphology (Ngoepe et al. 2023).

Many studies have investigated the genetic architecture and myriad signalling pathways underlying trophic morphologies in teleosts, and more specifically in cichlids (Albertson et al. 2005; Albertson and Kocher 2006; Parsons and Albertson 2009; Roberts et al. 2011; Parsons et al. 2014; Ahi 2016; Singh et al. 2021). In addition to gene expression variation, alternative splicing of pre-mRNA is also thought to contribute to transcriptional diversity and act as substrate for rapid adaptive evolution (Singh and Ahi 2022). Through alternative splicing a single gene can be altered into many mRNA isoforms which can potentially be translated into different proteins, boosting transcriptomic and proteomic diversity (Kim et al. 2007; Nilsen and Graveley 2010). Although it is debated to which extent alternative splicing events are contributing to proteomic and phenotypic diversity (Tress et al. 2017; Blencowe 2017). Splicing processes such as exonization, where exons evolve from retained introns, can also rapidly generate protein-coding and regulatory variation (Keren et al. 2010; Sorek 2007). It is generally thought that intron retention is rare in animals compared to plants, Singh et al (2017) found higher intron retention in cichlid fishes than humans. The relationship of alternative splicing for adaptive diversification and speciation is a research topic of increasing interest in animals and plants (Singh et al. 2017; Bush et al 2017; Smith et al. 2018; Smith et al. 2020; Wright et al. 2022; Rodríguez-Ramírez et al. 2023), but more research is needed to illuminate the role of alternative splicing during adaptive radiation, and more broadly adaptation and speciation (Singh and Ahi 2022).

Ontogenetic diet shifts, which are common in animals (Sánchez-Hernández et al. 2019), can alter the trajectory of adaptive radiations in the presence of ecological opportunity (Ten Brink and Seehausen 2022). Little attention has been directed towards the role of ontogenetic developmental changes during adaptive radiation (De-Kayne et al. 2024). In cichlids, there is some evidence that key shape changes in cichlid jaw cartilage and bone occur during early development (Albertson and Kocher 2006; Fujimura and Okada 2008) and that specialised morphologies are already distinct between trophic niches at the onset of exogenous feeding (Singh et al. 2017). But most transcriptional studies across multiple species have focused on one stage (Singh et al. 2017; Ahi et al. 2019; El Taher et al. 2020), and not much is known about the transcriptional patterns and mechanisms underlying the larval development of alternative trophic morphologies and further ontogenetic changes towards adulthood beyond a few model systems (Mazin et al. 2021).

Here we leveraged a comparative framework to investigate the differences and commonalities between two postembryonic key life stages in cichlid species: the end of the larval stage: stage 26 (Fujimura and Okada 2007) and the adult stage. The end of the larval development (stage 26) is marked by the complete absorption of the yolk sac into the body cavity and denotes the onset of independent foraging as the larvae are released from the mouthbrooding mother’s mouth. The adult stage (marked by the onset of breeding and the emergence of male nuptial coloration) stands for full maturity of the entire phenotype. To identify developmental genes and regulatory mechanisms involved in trophic differentiation, we explored the landscape of gene expression and alternative splicing underlying the trophic adaptations of the oral jaw apparatus (OJA) and pharyngeal jaw apparatus (PJA) (Fig. 1 C and D) in Lake Victoria cichlids (Fig. 1).

**Figure 1:**
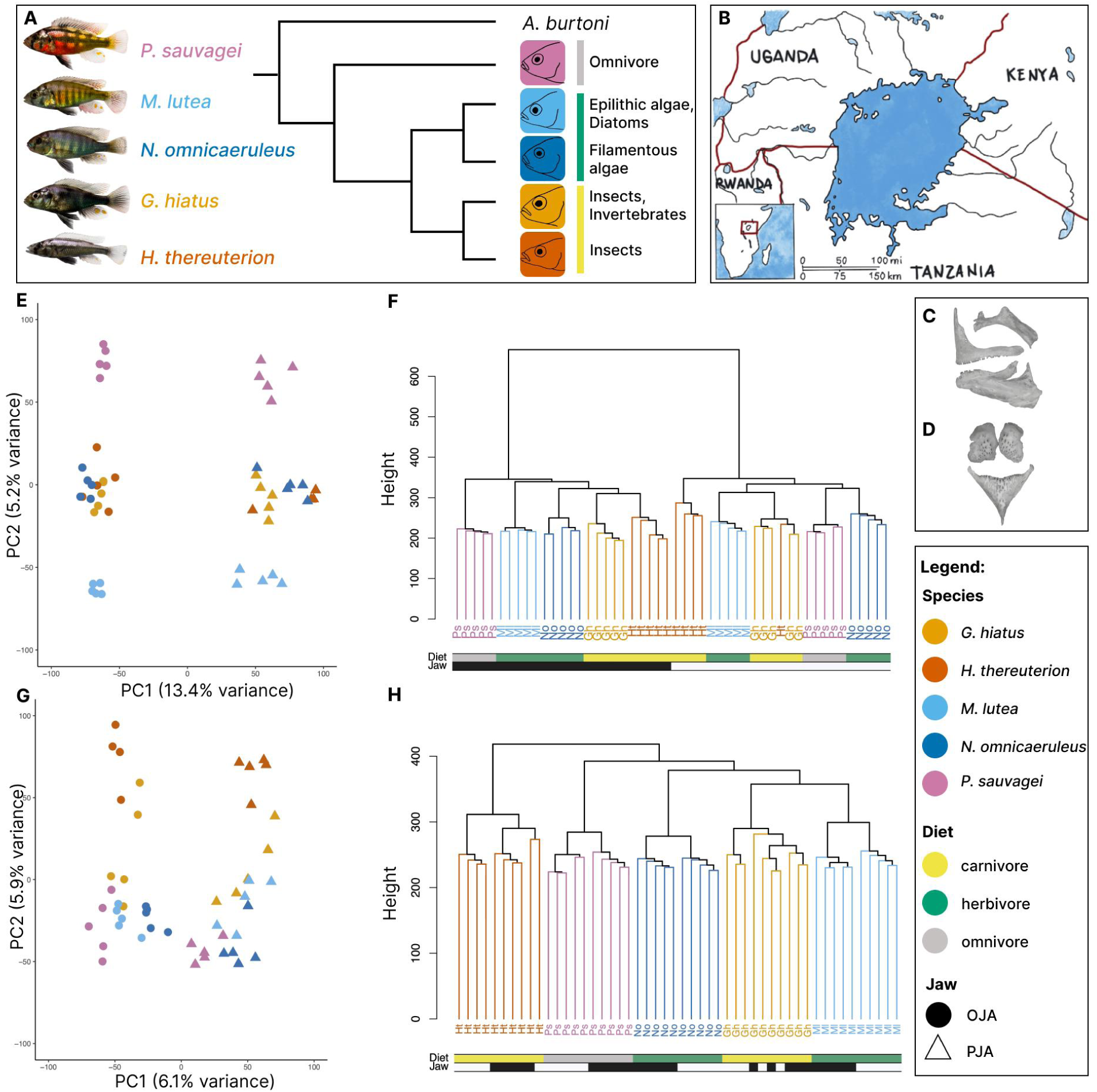
Overview of study design and broad patterns of gene expression. (A) Study species and their respective trophic niches. (B) Map of Lake Victoria. Photographs of Oral jaw apparatus (OJA) (C) and pharyngeal jaw apparatus (PJA) (D). Normalised gene expression PCA plot for larvae (E) and adults (G) based on 37,334 genes. (F) and (H) show dendrograms of the Euclidean distance based on the gene expression matrix in larvae (F) and adults (H). Oral jaw apparatus (OJA; black), pharyngeal jaw apparatus (PJA; white), H. thereuterion (Ht; dark orange), G. hiatus (Gh; bright orange), M. lutea (Ml; bright blue), N. omnicaeruleus (No; dark blue) and P. sauvagei (Ps; pink).

## Results

### Genome-guided transcriptome assembly and annotation

To analyse the transcriptional landscape of Lake Victoria (LV) cichlids adapted to different diets we generated whole mRNA transcriptomes from oral and pharyngeal jaws of five Lake Victoria species (*Haplochromis thereuterion* (Ht), *Gaurochromis hiatus* (Gh), *Mbipia lutea* (Ml), *Neochromis omnicaeruleus* (No) and *Paralabidochromis sauvagei* (Ps)) across two developmental stages (stage 26 - yolk sac absorption into body cavity and adulthood - onset of male nuptial colouration) (Fig. 1). Each stage and species was represented by five biological replicates. Tissues from two individuals were pooled for each biological replicate due to the small size of the larvae. Our total sample set comprised 100 whole transcriptomes (140 samples including the outgroup cichlid species *Tropheops tropheops* from Lake Malawi and an East African riverine species *Astatotilapia burtoni*). After quality filtering, we retained an average of 3-9 million reads per sample. Of these reads 68%-82% could be mapped to the reference genome of *O. niloticus* in the larval samples and 75%-82% in the adult samples (Supplementary File 1). We chose to map to *O. niloticus* because it is the best annotated reference genome and high quality transcript annotations were important for our analyses. The final merged annotation of all samples included 37,334 genes and the transcript annotation 59,849 transcripts.

### Transcriptome wide patterns of gene and isoform expression

We quantified the gene expression of 37,334 genes across all samples. To obtain an overview of global gene expression patterns we conducted a principal component analysis (PCA) based on normalised expression of 37,334 genes. The PCA shows a clear separation of expression profiles of the two jaw tissue sets for both life stages but the separation in jaws is less prominent in adults (Larvae: PC1 (13.4%), Adults: PC1 (6.1%))(Fig. 1 E and G). We also quantified expressions of 59,849 transcripts to see how isoform expression compared to the gene expression. The isoform expression PCA showed similar patterns as the larval gene expression profiles, with a clear distinction between the two jaw apparatuses but less differentiation among species (PC1 (11.4%)) (Fig. 2 A). Isoform expression clustering in adults is more species specific but less variation is explained by PC1 (5.6%) and PC2 (4.4%) when compared to the gene expression PCA (Fig. 2 C).

**Figure 2:**
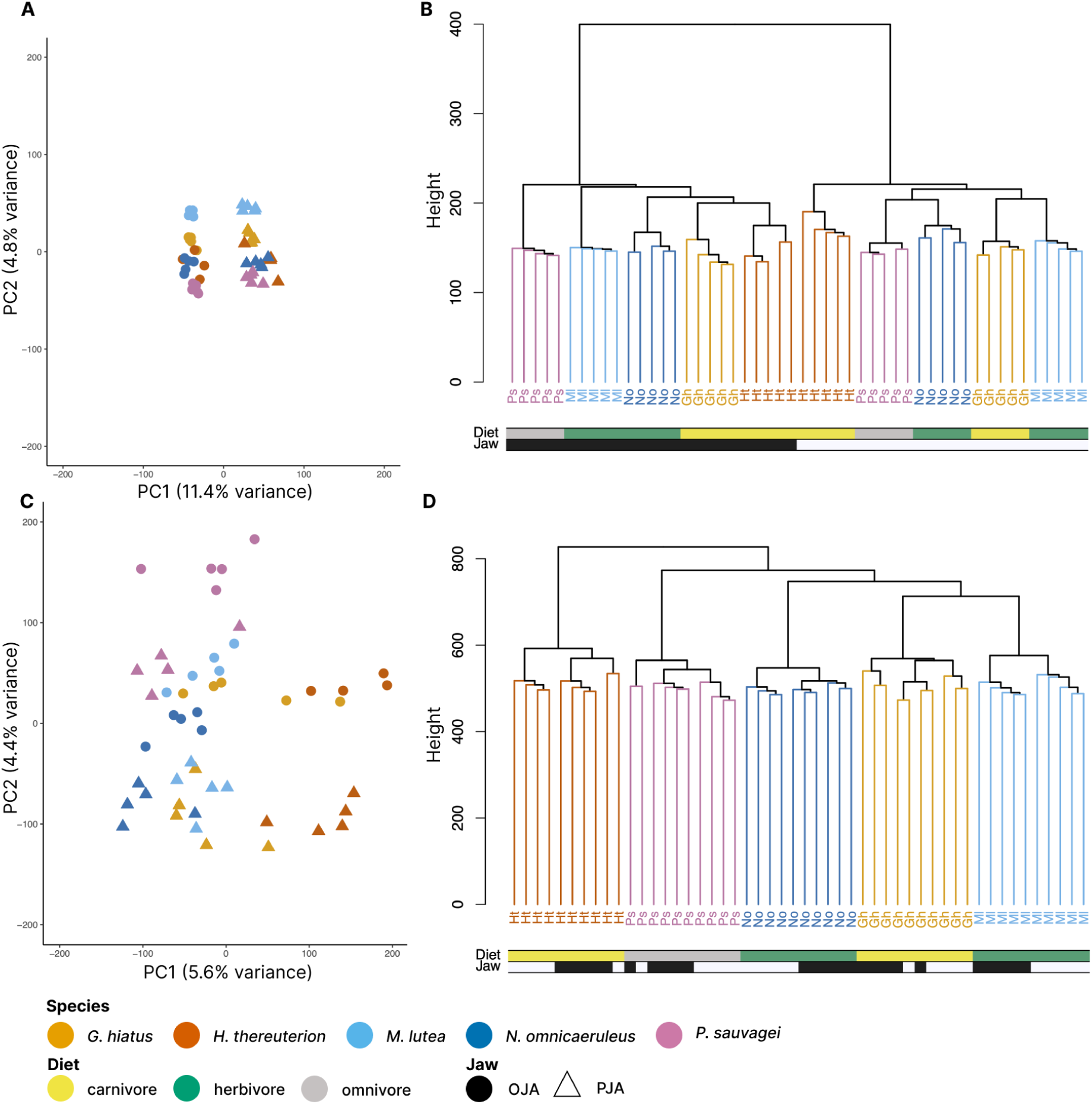
PCA plot for larvae (A) and adults (C) based on 59,849 normalised isoform expression counts. (B) and (D) show dendrograms of the Euclidean distance for larvae (B) and (D) adults. Oral jaw apparatus (OJA), pharyngeal jaw apparatus (PJA), H. thereuterion (Ht), G. hiatus (Gh), M. lutea (Ml), N. omnicaeruleus (No) and P. sauvagei (Ps).

Additionally we analysed the expression data using hierarchical clustering analysis (Fig. 1F-H). In the clustering analysis of the larvae, the expression of the two jaws is clearly distinct and the OJA clustering follows the trophic niche classification, while PJA clustering is species specific (Fig. 1 F). The clustering of gene expression in adults is strictly species-specific, regardless of jaw tissue (Fig. 1 H). The hierarchical cluster analysis of the isoform expression matrix shows a slight difference in the species patterning of the larval PJA compared to the hierarchical clustering of the gene expression matrix (Fig. 2 B), while the adult patterning stays the same (Fig. 2 D).

To place the gene expression patterns into a wider phylogenetic context, we included data from individuals of *Tropheops tropheops* (Tt) and *Astatotilapia burtoni* (Ab) at both stages as outgroups to extend the comparison to two more divergent species from Lake Malawi and the rivers around Lake Tanganyika respectively. The PCA as well as the hierarchical cluster analysis based on Euclidean distance shows a clear separation of expression profiles of *A. burtoni* and *T. tropheops*, from the Lake Victoria species (Fig. S1).

### Genes involved in species and trophic differentiation in Lake Victoria cichlids

To identify genes that contribute to species differences in oral and pharyngeal jaws of LV cichlids we conducted differential expression analysis for all pairwise species comparisons for each jaw and developmental stage. Of the 37,334 genes, 5,743 - 10,024 (15.4%-26.9%) showed significant (adjusted p-value < 0.05) differential expression (DE) between at least one pairwise species comparison in all tissues and at both developmental stages (Table S1). Between 591 - 1,664 (1.6% - 4.5%) genes were differentially expressed in more than 50% of the comparisons investigated. Interestingly, no gene was differentially expressed across all comparisons.

To identify genes that play a role in trophic adaptation of the jaw apparatuses we primarily focused on comparing the carnivorous species (Ht, Gh) to the herbivorous species (Ml, No). We identified the intersection set of genes that were differentially expressed across all four pairwise comparisons (Gh versus Ml, Gh versus No, Ht versus Ml, Ht versus No) (Fig. 3), and discovered 237 genes that were differentially expressed in larval OJA across all four herbivore versus carnivore comparisons and 159 genes were differentially expressed in the larval PJA of all four comparisons (Fig. 3 A and B respectively). In adult OJA comparisons we identified an overlap of 128 genes that were consistently differentially expressed and 107 genes were differentially expressed in the PJA (Fig. 3 C and D). Of these gene intersection sets between carnivores versus herbivores 79% - 96% shared the same direction of differential expression in all stages and jaw modules (Table S2). Such genes that were upregulated in carnivorous species and downregulated in herbivorous species or vice versa are interesting genes for further investigation in shaping divergent trophic morphologies.

**Figure 3:**
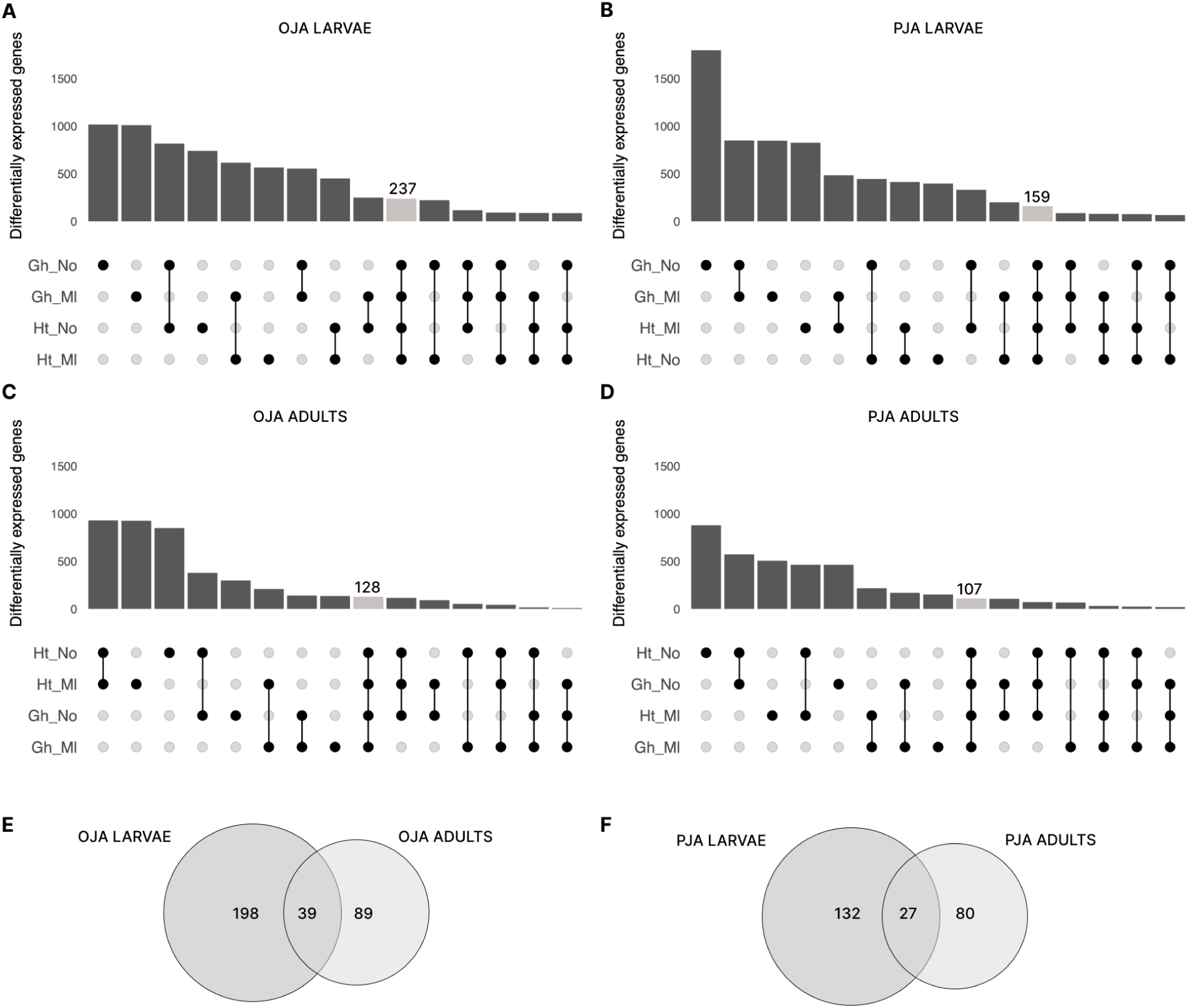
Differentially expressed genes between jaws of herbivorous and carnivorous Lake Victoria cichlids. Upset plots of overlapping differentially expressed genes for species comparisons including the carnivorous and herbivorous species. Points below bars show overlapping genes between indicated pairwise comparisons or unique sets of genes for single points. Light gray bars indicate an overlap between all pairwise comparisons. The total sum of differentially expressed genes for one pairwise comparison results from the sum of indicated bars. Upset plots are shown for larval OJA (A), larval PJA (B), adult OJA (C) and adult PJA (D). Venn diagrams show the overlap in the sets of shared differentially expressed genes for larvae and adults for OJA (E) and PJA (F). Oral jaw apparatus (OJA), pharyngeal jaw apparatus (PJA), H. thereuterion (Ht), G. hiatus (Gh), M. lutea (Ml) and N. omnicaeruleus (No).

We generated expression heatmaps with hierarchical clustering of all the aforementioned shared genes for each jaw tissue and developmental stage that were differentially expressed in all pairwise comparisons between carnivores and herbivores (Larvae: OJA: 273, PJA: 159; Adults: OJA: 128, PJA: 107; grey bars in Fig. 2 A - D). We also included the expression of these genes from omnivorous LV species *P. sauvagei*. The heatmap clustering placed *P. sauvagei* closer to the carnivorous *G. hiatus* and *H. thereuterion* for the oral and pharyngeal jaw gene expression at the larval (Fig. S2 A & B) and adult stages (Fig. S2 C and D). Additionally, we conducted Gene Ontology (GO) enrichment analysis to identify the biological processes that differentially expressed genes were involved in for these different stage and tissue comparisons (carnivorous versus herbivorous). We found that larval OJA genes were associated with jaw development related terms such as ‘roof of mouth development’ (*sox9b*) and ‘collagen catabolic process’ (*mmp2*); and also more general functions like ‘netrin-activated signalling pathway’ involved in axon extension and cell migration, ‘regulation of lipid catabolic process’ and ‘actin filament severing’ (Fig. 4 F, for full list see: Supplementary File 3). Genes differentially expressed in the larval PJA showed enrichment for GO terms involved in functions like ‘regulation of cell shape’ (*stat3*, *amotl2b*) and ‘skeletal muscle satellite cell migration’ (*rhocb*) (Supplementary File 3). In adult OJA, genes were significantly associated with the ‘the positive regulation of canonical wnt signalling pathway’ (*bambia*, *fam53b*) as well as ‘wnt signalling pathway’ (*bambia*, *bcl9l*, *fam53b*). Interestingly, genes differentially expressed in the adult PJA showed a significant enrichment in ‘regulation of alternative mRNA splicing, via spliceosome’ (*rbfox1*) (Full list in Supplementary File 3).

**Figure 4:**
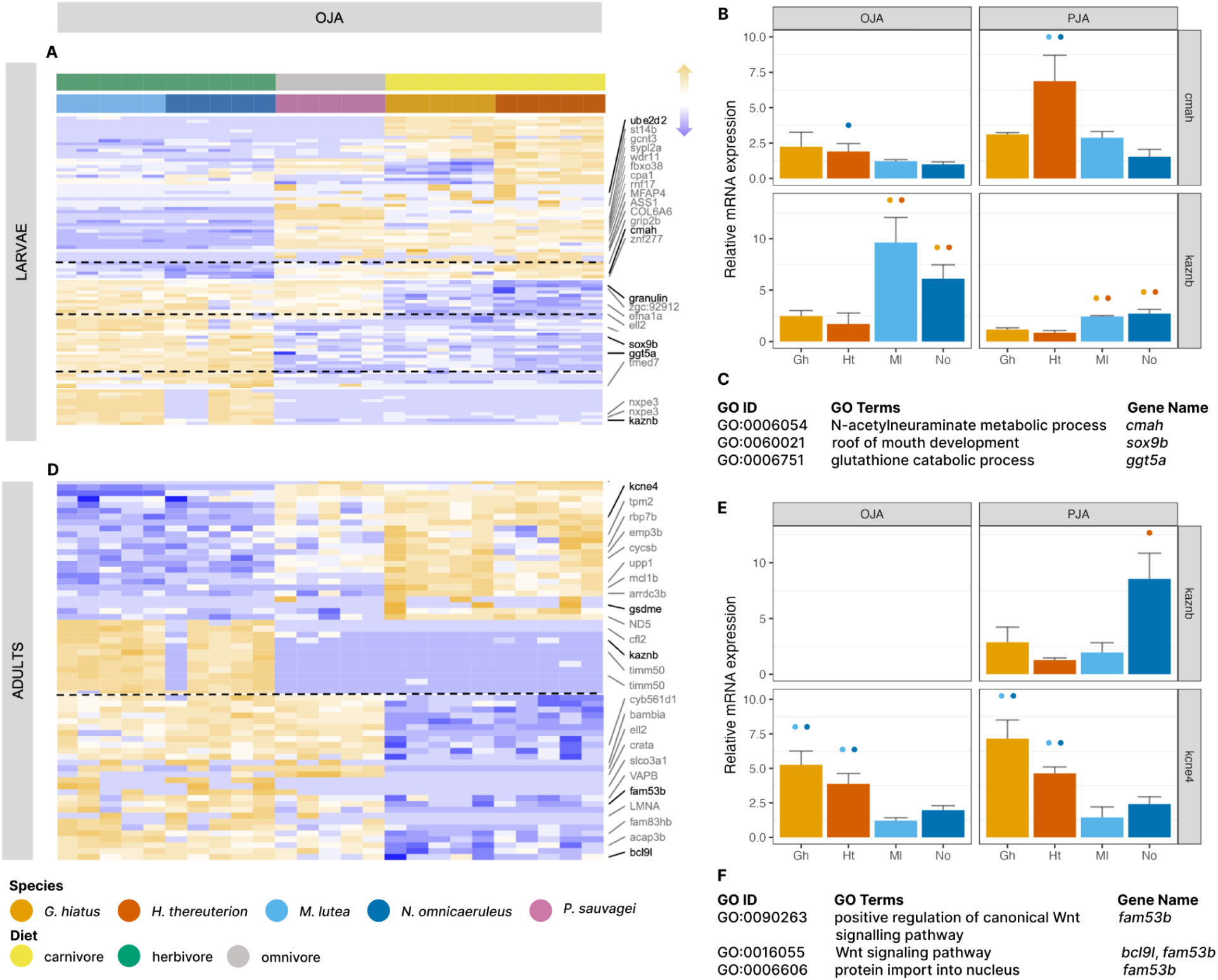
Genes involved in regulating herbivorous versus carnivorous jaw adaptations in Lake Victoria cichlids. Selected parts of heatmaps (dashed lines show cuts in heatmap) showing differentially expressed genes in comparisons including carnivorous versus herbivorous species for larval oral jaw apparatus (A) and adult oral jaw apparatus (D) respectively (Full heatmaps in the supplementary (Fig. S3 A - D)). Genes printed in black were either used in the qPCR validation analysis and/or have been significantly enriched for development and morphology related GO terms. Important jaw development associated GO terms are listed in (C) for larval OJA tissue and (F) for adult OJA tissue. The bar plots show qPCR validation of cmah and kaznb gene in larval (B) and kaznb and kcne4 in adult jaw tissues (E). Coloured dots above the bars indicate significant differences specified by species colours. Oral jaw apparatus (OJA), pharyngeal jaw apparatus (PJA), H. thereuterion (Ht), G. hiatus (Gh), M. lutea (Ml), N. omnicaeruleus (No) and P. sauvagei (Ps).

### Differentially expressed genes spanning both life stages

Comparing the intersection sets of differentially expressed genes between carnivorous and herbivorous species at both life stages showed an overlap of 39 genes in the OJA (25 downregulated and 13 upregulated in carnivores compared to herbivores between the following differentially expressed gene sets: larval OJA: 237; adult OJA: 128) (Fig. 3 E) and 27 genes in the PJA (18 downregulated and eight upregulated in carnivores compared to herbivores; between the following differentially expressed gene sets: larval PJA: 159; adult PJA: 107) (Fig. 3 F) (Supplementary File 2) across the life stages.

One gene that was consistently downregulated in the two carnivorous species as well as the omnivorous species in both tissues at both stages was *kaznb*, also called *kazrin* o*r periplakin-interacting protein b* (Fig. 4 A and D). We validated *kazbn* expression using qPCR analyses and found it mostly supports the RNA-seq results (Fig. 4 B and E). A more detailed inspection of the *kaznb* gene found that the differentially expressed transcript of interest is absent in the reference genome’s *kaznb* annotation. This transcript is spanning an intronic region in the *O. niloticus* reference and is not linked to any other annotated exons of *kaznb*. This observation suggests that exonization of introns is a mechanism through which both novel coding variation (new exon) and novel regulatory variation (new mRNA isoform) may have evolved in the ancestor of the of Lake Victoria cichlids or in the course of adaptive radiation within the lake.

### Gene expression modularity of the two jaws

Evolutionary modularity in gene regulation can contribute to morphological diversification and specialisation through the rewiring of gene regulatory networks (Wagner et al. 2007). We investigated gene expression differences between the oral and pharyngeal jaw apparatuses in herbivorous versus carnivorous species, to see whether the underlying gene regulatory architecture behind these functionally independent jaw modules differs between trophic niches. At stage 26 larvae, 1,889 (1768+121) genes were found to be differentially expressed between oral and pharyngeal jaw apparatus in all pairwise species comparisons, while at the adult stage only 209 (121+88) genes were consistently differentially expressed between the two jaws (Fig. 5 A; Fig. S3 A and B). One hundred and twenty-one of these genes were shared between both developmental stages, of which 68 were upregulated and 49 downregulated in the OJA versus PJA across all species and stages (Fig. 5 A). Four of these genes showed different directions of expression between the larval and the adult stage. Three of those four genes *ephb3*, *slc9a3* and *tbx3* were upregulated in larval PJA and downregulated in adult PJA. One gene, with the opposite expression pattern was unannotated. Several genes of the *Hox* family (*hoxa4a*, *hoxa4*, *hoxa5a*, *hoxb3a*, *hoxc5a*, *hoxc4a*) show a decreased expression in the OJA at both stages.

**Figure 5:**
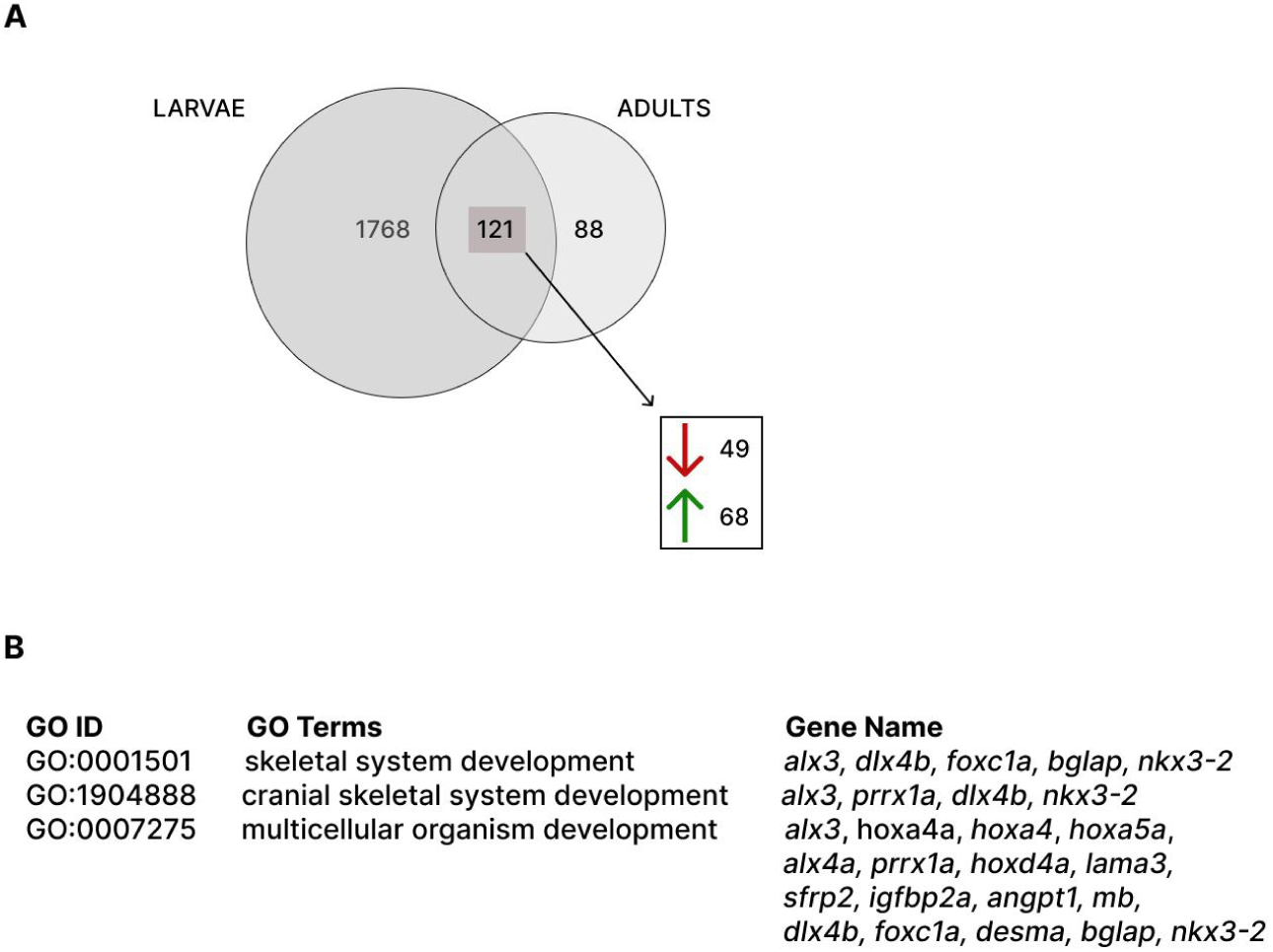
The Venn diagram shows the overlap in larval and adult differentially expressed genes in the oral versus the pharyngeal jaw apparatus and their direction of expression (upregulated genes are indicated by the upward facing arrow in green, downregulated genes by the downward facing arrow in red; direction indicated for OJA versus PJA) (A). Selected GO terms are listed in (B).

GO enrichment of the 121 overlapping differentially expressed genes between the OJA and the PJA of all carnivores and herbivores for both life stages showed significant enrichment for genes involved in ‘cranial skeletal system development’, regulation of muscle contraction’, ‘multicellular organism development’ and ‘regulation of transcription, DNA-templated’ (Fig. 5 B). Genes that were associated with the GO term ‘cranial skeletal system development’ include the genes *alx3, aristaless-like homeobox 3*, *prrx1a*, *paired related homeobox 1a*, *dlx4b*, *distal-less homeobox 4b* and *nkx3-2*, *NK3 homeobox 2* (Fig. 5 B).

### Quantitative PCR (qPCR) confirming gene expression

To validate our findings concerning differentially expressed genes between carnivores and herbivores we conducted qPCR analysis for five candidate genes for trophic divergence at the stage 26 larvae and four candidate genes for trophic divergence at the adult stage, which were found to be differentially expressed in both jaw apparatuses in our differential gene expression analysis. For the larval stage these included *cmah*, *ggt5a*, *granulin*, *kaznb* and *mpc1* and for the adult stage *kaznb*, *kcne4*, *mcl1b*, *slc5a8*. All of these differentially expressed genes showed the same direction of expression between the carnivores and the herbivores in our differential gene expression analysis. While not all of the qPCR differences were significant, they followed similar trends as our differential gene expression analysis results (Fig. S4). *cmah* was consistently upregulated in the differential gene expression analysis in both tissues for all carnivorous species (*G. hiatus and H. thereuterion*) at the larval stage (Fig. 4 A; Fig. S4 A), while the qPCR analysis for *cmah* in both larval tissues showed a significant difference in expression only between *N. omnicaeruleus* and *H. thereuterion* in the OJA and both herbivores (*M. lutea and N. omnicaeruleus*) compared to *H. thereuterion* in the PJA (Fig. 4 B). The differential expression analysis of *mpc1* showed an upregulation in larval herbivores (*M. lutea and N. omnicaeruleus*) in the PJA, which can be seen in *N. omnicaeruleus* in the qPCR analysis, but the difference is not significant (Fig. S4 A). The gene expression of *kaznb* was upregulated in herbivores versus carnivores in all tissues and all stages in the differential gene expression analysis. The qPCR analysis confirmed this pattern in both larval tissues, but at the adult stage, relative mRNA expression in the OJA of *kaznb* could not be detected, while in the PJA, a significant difference was detected only between *H. thereuterion* and *N. omnicaeruleus* (Fig. 4 E). The gene *slc5a8* did also not show significant differences between carnivorous and herbivorous species in relative mRNA expression in the adult OJA, despite it being upregulated in herbivorous species in the differential gene expression analysis (Fig. S4 B).

### Differential alternative splicing underlying trophic adaptation

To investigate the role of alternative splicing in trophic adaptation we conducted pairwise species comparisons between herbivorous and carnivorous species to detect differentially used isoforms. The numbers of differentially used isoforms ranged from 671 to 980 across all comparisons at each stage in both tissues (Table S3). A much smaller number of isoforms were significant across all pairwise comparisons (Fig. S5). In total we found 56 isoforms to be differentially used between carnivorous and herbivorous species in the OJA at the larval stage and 43 isoforms in the PJA at the larval stage (Fig. S5 A, B). At the adult stage we found 75 isoforms in the OJA to be differentially used and 78 isoforms in the PJA at the adult stage (Fig. S5 C, D). The intersection sets of differentially used isoforms between the stages per tissue reveals even less overlap with 20 isoforms between the OJA of larvae and adults and seven between the PJA (Fig. S5, E and F).

To visualise differential isoform usage, we chose the median PSI (percentage or proportion spliced in) values, describing the relative abundance of each isoform, for differentially spliced transcripts between our carnivorous and herbivorous comparisons (Fig. S6; Supplementary File 4). Two isoforms of *fgfr2* (fibroblast growth factor receptor 2) were differentially used in the OJA (Fig. 6 A & D) at both stages and the PJA of adults (Fig. S6) but showed a trophic signal only in the larval OJA tissue, with only *fgfr2.1* being expressed in the herbivores while both isoforms were used in the carnivores. In the adult OJA a gene called *frzb* (frizzled related protein) showed a clear distinction between the carnivorous and herbivorous species, whereby in the herbivores only one isoform is used (*frzb.1*), while in the adult carnivorous and the omnivorous OJA tissues the other isoform (*frzb.2*) shows the higher median PSI value (Fig. 6 E). *frzb* also is significantly differentially used in our comparisons of larval OJA tissue and showed up in the GO enrichment, however the pattern is more species-specific, with only *frzb.1* being abundant in larval Gh (carnivorous) and Ps (omnivorous) OJA tissue. The PJA of larvae showed the most distinct isoforms to be differentially used compared to the other three tissues (Fig. S6 B). Another gene that showed very similar patterns in the OJA tissues of both stages and the adult PJA is *col6a1*, collagen type VI alpha 1 chain, associated with maintenance of tissue integrity. The isoforms are more equally used in the carnivorous species, while in the herbivores *col6a1.1* shows a higher rate of usage than *col6a1.2* (Fig. 6 C & F).

**Figure 6:**
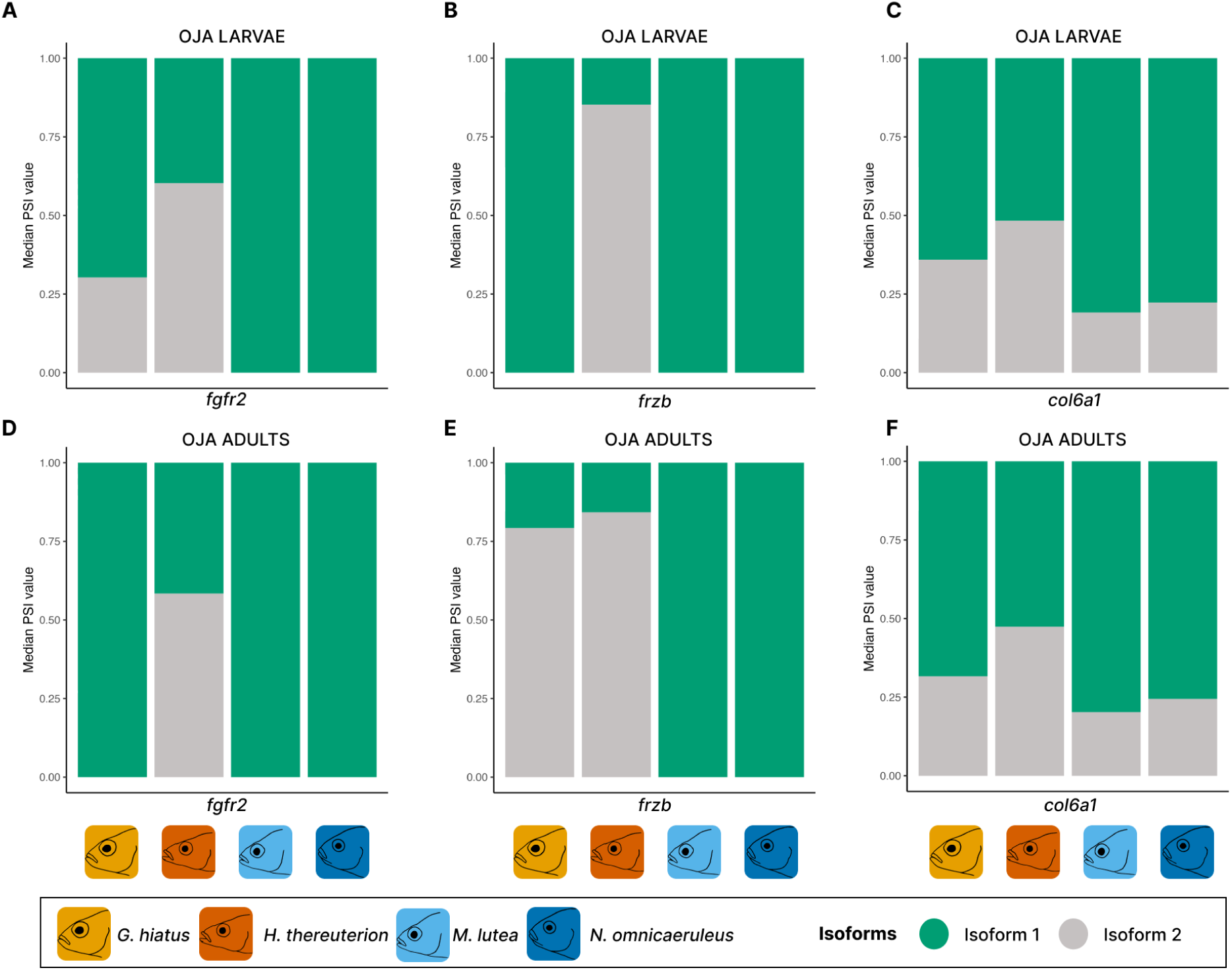
Median PSI values for isoforms, which were differentially used in carnivorous versus herbivorous comparisons in larval (A, B, C) or adult (D, E, F) OJA and PJA. Depicted are three selected genes (fgfr2, frzb and col6a1) and their respective median PSI values in the respective tissue. Oral jaw apparatus (OJA).

GO enrichment analysis of genes that were alternatively spliced showed *frzb* and *fgfr2* to be significantly enriched in the OJA at both stages, as well as the PJA of the adult stage (Supplementary File 5). *frzb*, was associated with ‘chondrocyte proliferation’ and *fgfr2* with ‘fibroblast growth factor receptor signalling pathway’ in larval and ‘chondrocyte proliferation’, ‘peptidyl-tyrosine phosphorylation’ and ‘peptidyl-tyrosine modification’ in the adult OJA as well as ‘positive regulation of cell proliferation’ and ‘transmembrane receptor protein tyrosine’ in adult PJA GO enrichment analysis.

### Contrasting roles for differential gene expression and alternative splicing

To investigate if genes that were differentially expressed were also alternatively spliced we looked at the intersection set between the two analyses. Interestingly, there was little to no overlap between differentially expressed and alternatively spliced genes between herbivores and carnivores (Fig. S7). Only one transcript was both, differentially expressed as well as differentially spliced in the OJA of larvae, but was not annotated with a gene name. In larval OJA, transcripts associated with three genes (*col6a1*, *efna1a* and *si:ch211-219a4.3*) were differentially spliced and differentially expressed. There was also little to no overlap in the enriched GO terms in differentially expressed and spliced genes (Fig. S8, Supplementary File 3 & 5). This suggests that gene expression and alternative splicing perform non-redundant functions.

## Discussion

Variation in gene regulation is more likely to be important for evolution at short timescales because new beneficial protein-coding and structural mutations take more time to evolve and establish. While the role of gene regulation has been studied in the context of adaptation and speciation (Mack and Nachman 2017, Singh et al. 2017, El Taher et al. 2020, Pavey et al. 2010), its developmental and ontogenetic dynamics are not well understood. The extremely young and ecologically diverse species flock of haplochromine cichlids from Lake Victoria in East Africa are a highly suitable model to test the relative importance of different levels of gene regulation during development across different species. Here we compared the gene regulatory landscape of Lake Victoria cichlid species adapted to divergent trophic niches at two post-embryonic life stages and found evidence for substantial developmental modularity in how genes are expression and spliced between species adapted to different trophic niches.

### Transcriptional dynamics of cichlid jaw development

Studying natural populations that have undergone adaptive radiation can provide insights into how organisms adapt to changing environments. It can help solve the evolutionary puzzle as to why some lineages diversify rapidly, while others do not, and which genetic and molecular mechanisms underlie these processes. We analysed the transcriptional landscape of oral and pharyngeal jaws in Lake Victoria cichlids and found a strong signal of gene and isoform expression conservation at the tissue level in larvae (Fig. 1 E & F, Fig. 2). Unlike the pharyngeal jaws that are used for processing food, oral jaw samples are clustered by trophic niche specialisation at the larval stage. Oral jaws are used to capture food and morphological differences in species-specific adult morphologies already arise during early juvenile development (from stage 26 onwards) (Fujimura and Okada 2008). This suggests that the gene regulatory pathways that are involved in trophic differentiation of oral jaws are active early in development, in preparation for when the larvae leave their mouthbrooding mother’s buccal cavity to start feeding independently. In contrast to the larvae, jaw gene and isoform expression in adults was species-specific (Fig. 1 G & H, Fig. 2). This means that the expression profile of oral and pharyngeal jaws of one species were more similar to each other than to the same tissue in a closely related species, so gene regulation is evolving rapidly in cichlid jaws. This is different to what has been observed in adult mammals where tissue-specific gene expression is conserved in deep evolutionary time of several million years (Merkin et al. 2012).

Developmental modularity is also reflected in the number of genes that are uniquely differentially expressed between oral and pharyngeal jaws at the larval vs adult stage (Figure 5). There are 1,768 genes that are only differentially expressed at the larval stage and 88 genes that are only expressed in adults, suggesting that these genes are performing unique functions during ontogeny - and that major differences in the two jaws are established early in development. Overall our results suggest that gene regulatory programs during jaw development are dynamic, controlling the basic development of jaw structures in larvae and later, in adults, shaping jaws to reflect species-specific trophic adaptations, presumably in response to environmental influences.

### Candidate genes for cichlid jaw modularity

It has been proposed that evolutionary decoupling (i.e. modularity) of jaws played an important role in cichlid diversification (Liem 1973). However, this hypothesis has mostly been tested in form and function (Liem 1973; Hulsey et al. 2006; Conith and Albertson 2021), or in gene regulation at one developmental stage (Singh et al 2021). Conith and Albertson (2021) emphasised the importance of studying integration/modularity at different taxonomic and developmental levels to better understand its role in cichlid adaptive radiation. As mentioned above, many genes that differentiated OJA versus PJA were unique to each developmental stage. However, a subset of genes were differentially expressed between the two jaws at *both* stages across all species (a total of 121 genes, which represented only 6,8% of the shared genes across species for larvae but 63.7% for adults) and showed the same direction of expression (except four genes) (Figure 5). This points to a conserved function of these 121 genes in differentiating oral vs pharyngeal jaws of these 121 genes at both life stages. Several *Hox* genes were part of this conserved gene set as *Hox* genes play an important role in the anteroposterior patterning of organisms and are known to not be present in the first pharyngeal arch, the mandibular arch (MA), forming the jaws in gnathostomes (Kuratani 2004; Ahi 2016). Recently it has also been shown that genes of the *Hox* family are not only expressed during development but also in adult tissues, contributing to bone regeneration (Rux and Wellik 2017; Bradaschia-Correa et al. 2019).

Interestingly, we found that *alx3*, which is known to be involved in cranial skeletal system development (Khor and Ettensohn 2020) was also in this developmental conserved gene set, being upregulated in the oral jaws at both life stages. In zebrafish (*Danio rerio*), *alx* genes are enriched in frontonasal neural crest cells (NCC) at early stages of development, playing a role in shaping skeletal elements of the anterior neurocranium (Mitchell et al. 2021). While *Alx3* has been repeatedly lost in chicken (*Gallus gallus*), a frog (*Xenopus tropicalis*) and lizard (*Anolis carolinensis*) species, its role might be compensated for by *Alx1* and/or *Alx4* (McGonnell et al. 2011). An alx1 haplotype has been associated with beak shape diversification across the adaptive radiation of Darwin’s finches (Lamichhaney et al. 2015). Similar to cichlid jaws, beak shape in finches has evolved in response to selection pressures to adapt to different food sources. Our findings suggest the ALX transcription factors may play a conserved function in craniofacial and trophic diversification during adaptive radiation across vertebrates.

### Repeated use of genes in the evolution of herbivory vs carnivory

Trophic diversification is a hallmark of adaptive radiation. While the transition from carnivory to herbivory seems limited to only some fish genera, including cichlids and carps, the repeated evolution of herbivory is even rarer but can be observed in modern haplochromines (He et al. 2015; McGee 2020; Singh et al. 2022). Unravelling the underpinnings of the evolution of herbivory and carnivory is key to understanding how trophic diversity shaped cichlid radiations. Singh et al. (2022) found that in Tropheini from Lake Tanganyika, that are closely related to the Lake Victoria cichlids, the molecular driver of repeated trophic transitions between herbivory and carnivory may be driven by regulatory variation rather than protein-coding mutations. As regulatory changes can evolve more rapidly than coding mutations, investigating the role of gene regulation is especially interesting in very young adaptive radiations.

To identify the genes involved in trophic adaptation we compared expression patterns in pairwise species comparisons between herbivorous and carnivorous species. A small number of shared differentially expressed genes across all species pairs indicated highly species-specific gene regulatory programs in Lake Victoria cichlids. The majority of genes (79% - 96%) that were differentially expressed between all carnivores versus herbivores pairs has the same direction of expression, suggesting a role of these genes in repeatedly contributing to trophic divergence. Several of these genes were associated with Wnt signalling, such as the *cmah* gene that was higher expressed in carnivores than herbivores at the larval stage (Fig. 4 A & B). *Cmah* encodes an enzyme that regulates Wnt signalling (Nystedt et al. 2009). The Wnt/β-catenin signalling pathway plays a crucial role in skeletogenesis of trophic structures, especially in remodelling and growth of fish jaws (reviewed in Ahi 2016). Higher expression of Wnt in Lake Malawi cichlids, was also correlated with a more steeply descending profile and early variation and increased ossification in craniofacial development (Parsons et al. 2014). Herbivory versus carnivory genes in OJA at the larval stage were enriched for GO terms like ‘roof of mouth development’ (Fig. 4 C) that was associated with genes *sox9b*, which is downregulated in the carnivorous species. *Sox9b* is a transcription factor involved in chondrogenesis and has been shown to be important for craniofacial development (Yan et al. 2005). The repression of *sox9b* expression through activation of the Ahr pathway resulted in the development of a shortened jaw (Xiong et al. 2008). The differential regulation of the Ahr pathway has been shown to play a key role in the trophic divergence of the OJA in Arctic charr (Ahi et al. 2015). In adults, *slc5a8* was upregulated in the jaws of herbivores. The role of *slc5a8* in skeletal development and morphogenesis is unclear, however, it has been shown to be a regulator of Wnt/β -catenin signalling in mammals (Hu et al. 2016). In the OJA of adults, significant GO enrichment terms included ‘wnt signalling pathway’ and ‘positive regulation of canonical Wnt signalling’ (Fig. 4 F) associated with the genes *fam53b* and *bcl9l*. *Fam53b* plays a pivotal role in regulating the Wnt pathway (Kizil et al. 2014); *bcl9l* has been linked to tooth development in mice (Cantù et al. 2017) and craniofacial malformations in zebrafish (Cantù et al. 2018). Both these genes showed a lower expression in carnivores versus the herbivores.

Besides genes related to the canonical Wnt/β-catenin signalling pathway we found *ggt5* to be downregulated in the jaw tissues of larval herbivores. *Ggt5* is a direct downstream target of *runx2*, a major osteogenic transcription factor (Tarkkonen et al. 2017). It is likely that increased expression of *ggt5* in skeletal tissue enhances bone resorption by induction of osteoclastogenesis which has also been observed in another member of the same gene family in mice (Hiramatsu et al. 2007). Thus, the lower expression of *ggt5a* in both jaws of herbivorous species may indicate differences in bone resorption level between the trophic niches. Interestingly, another gene with reduced expression in both jaws of herbivores, *granulin*, is also known for its key role in the stimulation of osteoclastogenesis through activation of RANK signalling (Oh et al. 2015). Taken together expression results for *ggt5a* and *granulin* may imply a distinct bone resorption process between carnivores and herbivores at the larval stage.

In adults, the most consistent expression differences in both jaws (validated by RNAseq and qPCR) between the trophic niches were observed for *kcne4* showing reduced expression in herbivores. The role of *kcne4* in skeletal development has remained unclear, however, it is known that *kcne4* directly interacts with calmodulin (CaM) (Ciampa et al. 2011), the ubiquitous Ca2+-transducing protein with an important role in jaw skeletogenesis (reviewed by Ahi 2016). The signal mediated by CaM has been already demonstrated as a key molecular mechanism contributing to variation in jaw structures among closely related species of birds and fish (Abzhanov et al. 2006; Parsons et al. 2009; Gunter et al. 2014). Interestingly, *kcne4* is identified as the only ion channel gene which showed an upregulated expression after mechanical loading in rats (Mantila Roosa et al. 2011) which is accompanied with increased bone formation. This could imply an increased bone formation in herbivores as has also been suggested by Albertson et al. (2012) when comparing the maxilla of a cichlid specialised in scraping tough, filamentous algae to a closely related more pelagic planktivorous species. The maxilla of the herbivore species is revealed to contain more bone and to be more resistant to bending, compared to the planktivorous species (Albertson et al. 2012).

### Alternative splicing contributed to trophic divergence of jaws

We still do not fully understand the role of alternative splicing in adaptive evolution, and how it interacts with gene expression regulation of genes. We found that similar to differentially expressed genes, alternatively spliced genes showed stage-specific and tissue-specific patterns. Some genes were alternatively spliced in both jaws at both developmental stages in herbivorous vs carnivorous species (Fig. 6). One of those genes that was significantly enriched for the GO biological process term ‘chondrocyte proliferation’ in oral jaws at both developmental stages (larval OJA and adult OJA) was *frzb*, frizzled related protein. While the two isoforms of this gene showed a species-specific usage pattern in the larval OJA, a trophic niche specific pattern in usage emerged in the adult OJA (Fig. 6 B & E). Kamel et al. (2013) showed that *frbz* plays an important role in craniofacial development and palate morphogenesis, with a knockdown of *frzb* in zebrafish causing the complete loss of the lower jaw. *frzb* also enhances the diffusion of Wnt by blocking short-range effects and increasing long-range gene responses (Rochard et al. 2016), making it a modulator of Wnt signaling and playing a role in the convergent-extension of the jaw skeleton, meaning that it might be an underlying molecular reason for the protrusion of the jaw in carnivores.

A gene with different isoform usage between the OJA of larval herbivores and carnivores was *fgfr2*, fibroblast growth factor receptor 2 (Fig. 6 A & D). Mutations in *fgfr2* have been associated with Crouzon syndrome, causing craniosynostosis (the premature fusion of sutures) and is also linked, among other features, to a short hypoplastic maxilla and mandibular prognathism (Reardon et al. 1994; Glaser et al. 2000), potentially linking differences in isoform usage to a steeper jaw profile in herbivores. In cichlids *fgfr2* has also been found to be involved in tooth development (Hulsey et al. 2016). Overall these results suggest that the alternative usage of spliced isoforms played a role in the rapid evolution of jaws adapted to different food sources in Lake Victoria cichlids. Alternative splicing has been shown to evolve faster than gene expression in deeply divergent vertebrates (Barbosa-Morais et al. 2012). Since the long evolutionary divergence time between those species is not comparable to the very young Lake Victoria cichlids studied here, alternative splicing seems to already play an important role in formation of adaptive morphologies at very early stages of evolutionary divergence.

### Exonization generated novel variation for the evolution of herbivory

Exonization, the process of non-coding intronic regions becoming protein-coding exons, is a mechanism of genome evolution that can generate new gene isoforms (Sorek 2007). Although most exonization events are believed to be molecular noise, it provides a rapid mechanism for generating new evolutionary solutions from the original genome. As long as the original isoform maintains functionality, the new alternative isoform can avoid purging through selection and may evolve a novel function over time (Schmitz and Brosius 2011). Because exonization can rapidly evolve from standing variation, it has the potential to be an important generator of adaptive isoforms during evolution.

When we compared the jaw transcriptomes of herbivorous and carnivorous cichlids in our study, we found that only a small fraction of genes were differentially expressed between all carnivores and herbivores across both stages within each jaw and just a few of those were annotated (Supplementary File 2). One of these was a transcript of *kaznb*, also called Kazrin, encodes periplakin interacting protein b, and is involved in several cellular functions such as cytoskeletal organisation, epidermal differentiation, and cell adhesion (e.g. Sevilla et al. 2008; Cho et al. 2011). The expressed transcript falls into an intronic region in the *O. niloticus* reference and was consistently expressed in the same direction across all tissues and stages with lower expression in the carnivorous (and the omnivorous) species and higher expression in herbivores. We were able to validate the expression profile of the *kaznb* gene in the larval tissues via qPCR analyses (Fig. 4 B) and to some extent in the pharyngeal jaws of the adult tissues (Fig. 4 E). A study in *Xenopus laevis* larvae concentrating on the function of *kaznb* showed that it plays an essential role in craniofacial development, through controlling neural crest cell (NCC) establishment (Cho et al. 2011). When depleting *X. laevis* embryos for *kaznb* in the developing head region with morpholino oligo injection that blocked *kaznb* translation, Cho et al. (2011) detected eye and head defects, with smaller ceratohyal and ceratobranchial cartilages and eyes on the treated sides, suggesting an important role in craniofacial development. This seems to be an interesting candidate gene that has not been suggested so far to be relevant in shaping divergent cichlid trophic morphologies to our knowledge. Future studies should address its function in differences between herbivorous and carnivorous jaw structures.

It has been previously shown that specialised herbivorous and carnivorous haplochromine cichlids had significantly higher intron retention in their mRNA than omnivorous species (Singh et al. 2017). Some retained introns may represent exonized introns that are translated produce a novel protein with new domains. As exonization provides a relatively rapid way to generate protein coding variation and potentially evolutionary novelty, it could play a underappreciated role in adaptive radiation.

### Distinct roles of gene expression and alternative splicing during adaptive radiation?

A number of studies have empirically investigated the differing roles of differential gene expression and alternative splicing in phenotypic adaptation and found them to play non-redundant roles (Jakšić and Schlötterer 2016, Jacobs and Elmer 2021). In the butterfly *Bicyclus anynana* alternative splicing controlled a smaller set of genes than differential splicing, but these genes had unique adaptive functions (Steward et al. 2022). This is similar to what was found in cichlid fishes from Lake Tanganyika where differentially spliced genes were involved in adaptive trophic morphologies and differentially expressed genes were largely involved in housekeeping functions (Singh et al. 2017). In contrast, a recent study on the Eda haplotype in sticklebacks found only a small overlap (six genes) of gene expression and alternative splicing, but a lot of differentially spliced genes (no significant enrichment) were involved in the same biological processes as the differentially expressed ones (Rodríguez-Ramírez et al. 2023). In our study, almost no genes (or enriched GO biological processes) were both differentially expressed and differentially spliced between species adapted to different trophic niches (Fig. S7, Fig. S8). Even though these two gene regulatory mechanisms are targeting different genes, these genes are often involved in pathways related to the same broader craniofacial developmental process. For example, the alternative splicing GO enrichment showed involvement in ‘chondrocyte proliferation’ (Larval OJA, Adult OJA and Adult PJA) while the differential expression GO enrichment was associated with terms like ‘roof of mouth development’ (Larval OJA) and ‘Wnt signalling pathway’ (Adult OJA). Interestingly, one of the genes that was was both differentially expressed and alternatively spliced was col6a1, a gene implicated in differentiating cichlid oral and pharyngeal jaws in other haplochromine cichlids (Singh et al. 2017). This gene controls the mechanical properties of the connective tissue between muscle and bone, and thus may require complex regulation through multiple mechanisms. Even though the community is eager to identify broad patterns, based on what is currently known, it is not possible to predict which genes will be regulated by alternative splicing or gene expression. A meta-analysis of gene expression and alternative splicing across the tree of life could help shed light on this issue.

## Conclusion

Little is known about the developmental dynamics of gene expression and alternative splicing evolve at very short time scales as most studies published so far have focused on older lineages and single life stages. Here we show that both differential expression and differential splicing are important regulatory processes for driving rapid morphological change during development of divergent adaptations. Future studies would benefit from including multiple developmental stages in their study design and a broader phylogenetic sampling. Furthermore, we find that exonization may be a notable evolutionary mechanism for generating novel protein variation during rapid adaptive radiation. However, more research is needed to determine how functional and widespread such events are.

## Material and Methods

### Sampling

In this study we focus on five model species from Lake Victoria (*Gaurochromis hiatus* (*Gh*), *Haplochromis thereuterion* (*Ht*), *Mbipia lutea* (*Ml*), *Neochromis omnicaeruleus* (*No*) and *Paralabidochromis sauvagei* (*Ps*)). We picked those five species to try to reflect the broad spectrum of trophic specialisations in Lake Victoria. *G. hiatus* feeds on chironomid larvae as well as on bivalves and mollusks (Witte et al. 1982). *H. thereuterion* mostly feeds on insects like chironomid larvae, but a substantial portion of its nutrition is covered by collecting terrestrial insects from the water surface (Oijen et al. 1996). They both represent differentially specialised carnivores. *M. lutea* and *N. omnicaeruleus* are both mainly herbivorous but with different modes of feeding. While *M. lutea* feeds mainly on filamentous algae and also takes in some detritus, *N. omnicaeruleus* grazes off epilithic algae and diatoms (Seehausen et al. 1998). *P. sauvagei* is an omnivore, feeding on epilithic algae, associated insects and sponges (Seehausen 1996). As an outgroup we included *Tropheops tropheops* (Tt), an herbivorous Lake Malawi cichlid, which mainly feeds on algae and scratches it from hard surfaces, but has also been reported to exclusively feed on plankton if abundant (Konings 1990). As a second outgroup species we selected *Astatotilapia burtoni* (Ab), a mostly riverine cichlid that is located in Lake Tanganyika and its surrounding waterways and can be classified as an omnivore, feeding on plants, aufwuchs, insects and fish remains (Muschick et al. 2012). We scanned the pharyngeal jaw apparatus of *H. thereuterion*, *N. omnicaeruleus* and *P. sauvagei* at the larval and the adult stage to visualise the differences in morphology that are visible from the beginning of the exogenous feeding (Fig. S9). Scans were conducted with a Scanco Medical μCT 40 and images were created with the 3D Slicer version 5.6.2.

The *M. lutea* were collected from Makobe Island and the *G. hiatus* from the northern Mwanza Gulf. Both were collected by Ole Seehausen. The other samples, *N. omnicaeruleus*, *P. sauvagei* and *H. thereuterion* were obtained via aquarium trade. All fish were reared in the fish facility at University of Graz under standardised aquarium conditions regarding tank environment, diet and water parameters. Eggs were taken from the mouth of their mother 24 hours after fertilisation, by applying slight pressure onto their cheeks, and reared in a separate tank until the yolk sac was completely absorbed (stage 26, (Fujimura and Okada 2007)). The larvae were euthanized with an overdose of MS 222 and stored in RNAlater in a fridge for seven days and then moved to a -80°C freezer. Fish raised to the adult life stage were fed with a standardised diet of green spirulina flakes. When individuals started to show adult colouring and breeding behavior males were sacrificed with MS 222 and dissected. Bones were stored in RNAlater in a fridge for seven days and then frozen at -80°C.

### RNA extraction

Per species and stage five biological replicates of oral and pharyngeal jaw apparatus samples were taken. The oral jaw apparatus samples contained the premaxilla, maxilla, dentary and articulate while the pharyngeal jaw apparatus samples were composed of the upper and lower pharyngeal jaw. In the case of the larvae the area of the OJA and of the PJA were sampled under a stereo microscope and submerged in a lysis buffer and 1-thioglycerol according to the protocol of the Promega RNA Reliaprep kit. This kit includes filter columns and a DNA digestion step. The sample in the lysis buffer was crushed with a ceramic bullet in a homogenizer (FastPrep-24, MP Biomedicals, Santa Ana, CA, USA), before further processing. Two individuals were pooled per sample to increase the amount of RNA. For the adult species the bones were all cleaned from skin and muscle tissue and stored in RNAlater. As the whole set of bones would have been too much tissue input for the RNA extraction, only half of the bones were collected. The set of the bones is the same for all samples, but if bones for RNA extraction were collected from the left or right side was randomly decided by a simple randomization script in R (R Core Team 2020) prior to each extraction. RNA for the adults was extracted using TRIzol reagent (Thermo Fisher Scientific), following the protocol of the TRIzol protocol for RNA extraction with the inclusion of a DNAse step at the end (invitrogen). RNA quality was tested at a TapeStation 2200 with RNA Screen Tapes. It was tried to reach a RINe number of >= 7 for each sample.

### Library preparation and sequencing

Library preparation was done with the TruSeq Stranded mRNA Sample Prep Kit using 24 indexing adapters and following the standard protocol. For the input RNA we tried to reach 1000 ng for each sample. The quality of the cDNA was tested on a TapeStation 2200 (Agilent) with D1000 Screen Tapes from the Agilent RK6 ScreenTape System. Sequencing was executed at the Vienna BioCenter Core Facilities on a HiSeq 2500 with paired end 125 cycles per read (2 x 125 cycles). Demultiplexing was conducted by the same facility.

### Transcriptome assembly

For each of the samples we got an average of ten million paired-end reads. Sequence reads are available at NCBI sequence read archive (SRA) under the accession number PRJNA640176. After a quality check with Fastqc (v0.11.8) (FastQC 2015) and a trimming step with Trimmomatic (v0.3.9) (Bolger et al. 2014), only reads with a phred > 28 and a minimum length of 70 bp were retained. For each sample a median of about 5 million reads were obtained. For read alignment we used STAR(v2.7.3.a) (Dobin et al. 2013) with the *Oreochromis niloticus* genome reference from the University of Maryland (Conte et al. 2017) (*O. niloticus*; NCBI accession number: GCA_001858045.3) as guide reference, as it is very well annotated and this is important for accurate isoform reconstruction. After checking the mapping statistics with samtools idxstats (v1.9) (Danecek et al. 2021) and merging the single files for the same species and jaw type with picard (v2.21.7) (Picard toolkit 2019) we used StringTie (v2.0.6) (Pertea et al. 2015) without a reference to assemble the RNA-seq alignments into potential transcripts. This was done separately for the single files (per fish) and the merged files (per species and jaw). The single files for each biological replicate were then merged according to species, to subsequently merge all the files per species, tissue and stage. These repeated merging steps were conducted to reduce the probability of false positives. To estimate the accuracy of the produced annotation files we compared them with gffcompare (v0.11.2) (Pertea and Pertea 2020) to our reference annotation. We filtered for monoexonic transcripts not contained in the reference and a class code assigned by gffcompare, indicating ‘possible polymerase run-on’ fragments. The maximum intron length was reduced to 200000 bp, which is the maximum intron length found in the *O.niloticus* reference. Based on these annotations, the expression estimates were generated with StringTie allowing no multimapping. From these expression estimates, count matrices were produced using the prepDE.py script that comes with StringTie (v2.0.6) to extract raw count data from StringTie results. The code used for the whole analysis is stored in a github repository (https://github.com/annaduenser/LV_RNAseq).

### Differential gene expression analysis

We were not able to directly compare jaws in adults and larvae as we were not able to apply an identical dissection and extraction process at both life stages due to the minimal size of the larvae. We counteracted this shortcoming with not directly comparing the data sets, but conducting the same species-pairwise comparisons and further investigating the intersection set between those. This method should reveal identical patterns in gene expression and alternative splicing between trophic niches, if present. When looking at the Euclidean cluster dendrogram at the larval stage, that also includes our outgroup, we see a clear distinction between the LV species and the latter. When including species from Lake Malawi (LM) and Lake Tanganyika (LT), a jaw tissue specific pattern is still visible for LV as well as for LM (clustering next to LV), but not for LT (Singh in preparation) suggesting that this separation is not solely caused by the inclusion of surrounding muscle tissue. If anything, the surrounding tissue could potentially add to the trophic niche clustering in the larval oral jaw apparatus. We used DESeq2 (Love et al. 2014) in (R Core Team 2020) using RStudio (RStudio Team 2019) to detect differentially expressed genes running comparisons between all species and tissues as well as comparisons between the tissues within a species. DESeq2 uses raw read counts and estimates variance-mean dependence based on a model that utilizes the negative binomial distribution (Love et al. 2014). The cutoff for differentially expressed genes was chosen at a false discovery rate of (p < 0.05).

### GO term enrichment

GO terms for *Oreochromis niloticus* were acquired in R (R Core Team 2020) using RStudio (RStudio Team 2019) via the biomaRt package (v2.46.1)(Durinck et al. 2005; Durinck et al. 2009) in R from the Ensembl database. Gene set enrichment analysis was conducted with the R program topGO (v2.36.0)(Alexa and Rahnenfuhrer 2019) using the method *weight* to account for GO topology and a Fisher’s exact test for the enrichment analysis. We used our annotation to build the gene universe, where we then tested the list of genes, overlapping between carnivorous and herbivorous species in gene expression or alternative splicing analysis for gene set enrichment.

### qPCR analysis

From four replicates of RNA samples of each jaw apparatus, stage and species, we used 500 ng of RNA to synthesise first strand cDNA using the High Capacity cDNA Reverse Transcription kit (Applied Biosystems). The cDNA from each sample was diluted 1:10 times in nuclease-free water to be used for qPCR. For the adult stage, we selected five candidate target genes; *kcne4*, *kaznb*, *mcl1b*, *rnf114* and *slc5a8*, and for the larval stage we also chose five genes; *ggt5a*, *kaznb*, *granulin*, *faah* and *cmah* (Fig. S5). We used *rpl18* as a reference gene, which we have already validated as a stably expressed gene in both cichlid jaw tissues across different trophic niches (Ahi et al. 2019), to normalise our expression data for qPCR. The primers were designed at conserved sequence regions of the coding sequences from all species using CLC Genomic Workbench, version 7.5 (CLC Bio, Aarhus, Denmark) for alignment and delineation of exon boundaries using *A. burtoni* annotated genome in the Ensembl database (<http://www.ensembl.org>) (Zerbino et al. 2018). Primer Express 3.0 (Applied Biosystems, CA, USA) and OligoAnalyzer 3.1 (Integrated DNA Technology) were used to design the primers as described in details in our previous studies (Ahi et al. 2018; Ahi et al. 2019) (Supplementary data for primer info). Maxima SYBR Green/ROX qPCR Master Mix (2X) (Thermo Fisher Scientific, Germany) was used to generate qPCR reactions. The amplification steps were conducted in 96 well-PCR plates on ABI 7500 real-time PCR System (Applied Biosystems). The primer efficiency calculation, qPCR program, plate set-up and a dissociation step were performed as described previously (Ahi et al. 2018; Ahi et al. 2019). To normalise the expression levels, the Cq value of *rpl18* was used as normalisation factor (Cq reference), and Cq of each target gene was calculated accordingly (ΔCq target = Cq target - Cq reference). Relative expression quantities (RQ) were calculated for the normalised values using E-ΔCq (Pfaffl 2001) and then fold difference values were calculated by transformation of RQ values to logarithmic values in order to conduct further statistical analysis (Bergkvist et al. 2010). The statistical expression differences were identified using ANOVA statistical tests, followed by Tukey’s HSD post hoc tests.

### Alternative splicing analysis

As the alternatively spliced isoforms can be a result of different types of splicing, it is not as straightforward to assess but can be explored from different angles. Either from an exon centric approach via differential exon usage, or with an isoform centric approach, which can be subdivided by differential transcript expression and differential transcript usage. We used the approach of differential transcript usage analysis in this study to investigate differences in transcript usage between the different species and trophic niches. Alternative splicing analysis was conducted with SUPPA2 (v2.3) (Trincado et al. 2018) to evaluate differential transcript usage between the samples. SUPPA2 takes the normalised isoform counts of all replicates of a condition and calculates a relative abundance for each transcript in each sample (PSI). To detect the difference in transcript usage between conditions dPSI is computed (dPSI = PSI condition 2 - PSI condition 1) (Trincado et al. 2018). By taking the dPSI comparisons of herbivore versus carnivore species and selecting the isoforms that were significantly used differently. We illustrated the difference in usage by computing the median PSI value over the five biological replicates for each species for each transcript. Visualisation of the analysis was conducted in R (R Core Team 2020) using RStudio (RStudio Team 2019).

## Data availability

Sequence reads are available at NCBI sequence read archive (SRA) under the accession number PRJNA640176 (). The code for the analysis is available at the following github repository (https://github.com/annaduenser/LV_RNAseq).

## Acknowledgments

The authors acknowledge the financial support by the University of Graz. This work was supported by the Austrian Science Fund (project number P29838) awarded to CS.

